# Direct and indirect effects of land-use intensity on plant communities across elevation in semi-natural grasslands

**DOI:** 10.1101/2020.03.18.996744

**Authors:** Oksana Y. Buzhdygan, Britta Tietjen, Svitlana S. Rudenko, Volodymyr A. Nikorych, Jana S. Petermann

## Abstract

Grassland biodiversity is among the most vulnerable to land use. How to best manage semi-natural grasslands for maintaining biodiversity is still unclear in many cases because processes may depend on environmental conditions and indirect effects are rarely considered. Here we evaluate the relative importance of direct and indirect effects of grazing intensity on plant communities along an elevational gradient on a large topographic scale in the Eastern Carpathians in Ukraine. We sampled 31 semi-natural grasslands exposed to cattle grazing in two years. Within each grassland site we measured plant community properties such as the number of species, functional groups, and the proportion of undesirable weeds. In addition, we recorded cattle density (as proxy for grazing intensity), soil properties (bare soil exposure, soil organic carbon, and soil pH) and densities of soil decomposers (earthworms and soil microorganisms). We used structural equation modelling to explore direct and indirect effects of grazing intensity on plant communities along the elevation gradient. We found that cattle density decreased plant species and functional diversity but increased the proportion of undesirable weeds. Some of these effects were directly linked to grazing intensity (i.e., species richness), while others (i.e., functional diversity and proportion of undesirable weeds) were mediated via bare soil exposure. Although grazing intensity decreased with elevation, the effects of grazing on the plant community did not change along the elevation gradient. Generally, elevation had a strong positive direct effect on plant species richness as well as a negative indirect effect, mediated via altered soil acidity and decreased decomposer density. Our results indicate that plant diversity and composition are controlled by the complex interplay among grazing intensity and changing environmental conditions along elevation. Furthermore, we found lower soil pH, organic carbon and decomposer density with elevation, indicating that the effects of grazing on soil and related ecosystem functions and services in semi-natural grasslands may be more pronounced with elevation. This demonstrates that we need to account for environmental gradients when attempting to generalize effects of land-use intensity on biodiversity.

## Introduction

Grasslands cover more than 40% of the global terrestrial area [1] and are among the most species-rich habitats in Europe [2]. As a result of their high biodiversity, grasslands provide crucial ecosystem functions and services beyond that of livestock forage production [3–7]. However, grassland biodiversity compared to the diversity in other ecosystem types is among the most vulnerable to human impact, particularly to land-use change [8]. Of extraordinary importance for biodiversity are semi-natural grasslands [2,6,7], which are remnants of habitats created by tree cutting, haymaking, or low-intensity traditional farming [5]. In order to survive and to function, semi-natural grassland communities require regular grass removal, for example through grazing [2]. However, the effects of grazing on grassland biodiversity have been found to depend on land-use intensity [2,9] and on particular environmental conditions [10], yielding contrasting results. The inconsistency in the observed patterns can depend upon the balance of different mechanisms underlying the relationships among grazing and plant community composition and diversity along environmental gradients (summarized in Table S1 and Fig. 1).

**Fig.1.**
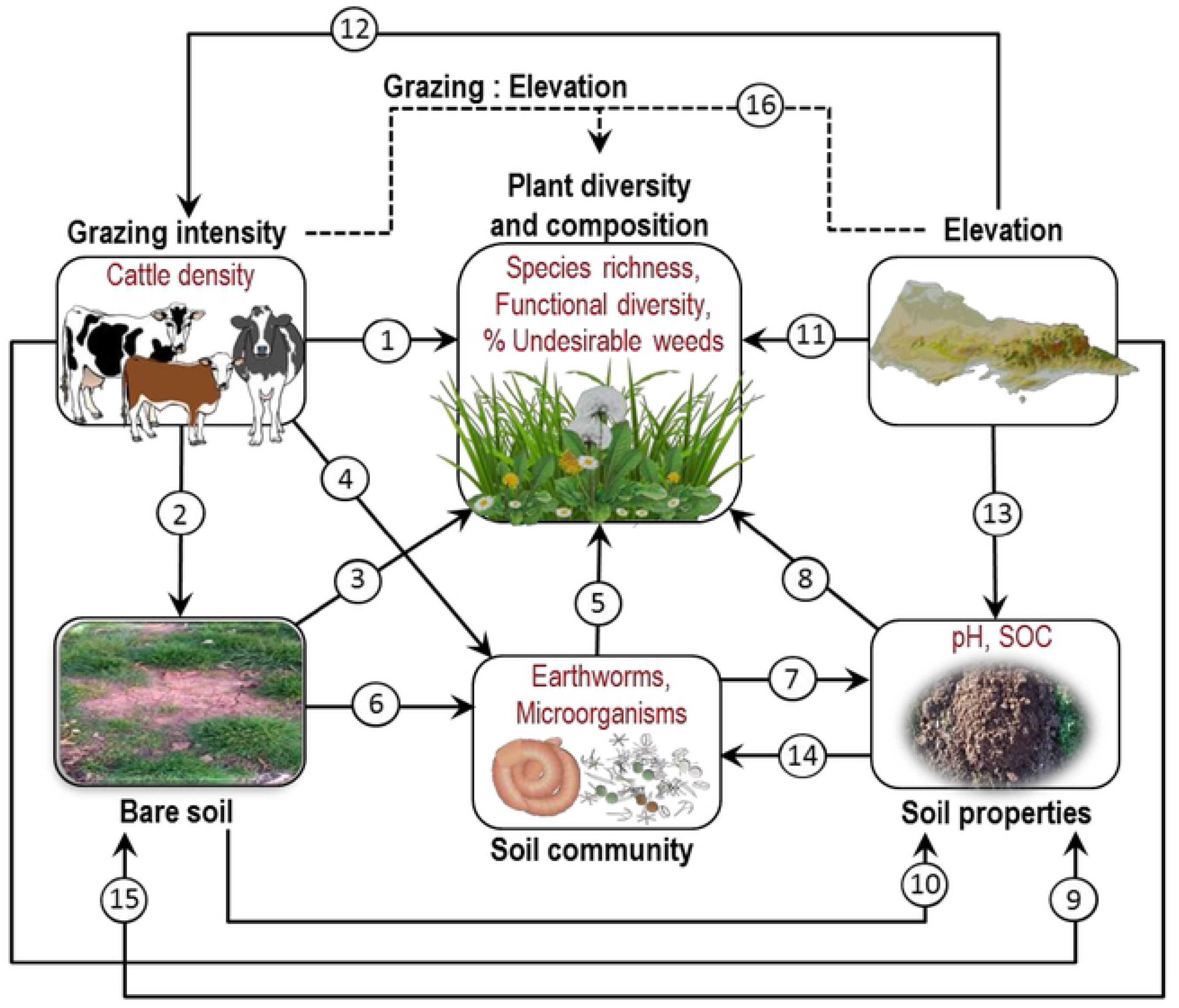
Conceptual model of the expected causal direct and indirect effects of grazing intensity and environmental conditions on plant diversity and composition. The conceptual model is based on hypotheses derived from previous research findings (see Table S1 for hypotheses and references). SOC, soil organic carbon. Dashed path 16 shows an interactive effect of grazing and elevation. The images were created using the IAN Symbols [59]. Photos of soil and bare soil: O. Buzhdygan.

The direct impact of cattle on a plant community (Fig. 1, *path 1*) falls into several different categories [11], and includes mechanical, chemical [12], and biological [6] effects on vegetation (Table S1). Grazing can impede as well as improve local colonization processes by plants for example through reduction of propagules of extant species or by increasing the dispersal of propagules of new species to a site [13]. Additionally, grazing can increase plant diversity by reducing competition via direct consumption of competitively dominant plant species [10]. By contrast, if grazing promotes species dominance by increasing the abundance of grazing–resistant, unpalatable species, then resource availability for other plant species decreases, thus reducing biodiversity [10]. In parallel, grazing can increase the spatial heterogeneity of vegetation [14], in comparison to e.g. hay meadows, which in turn could benefit plant diversity.

Effects of cattle grazing on the plant community can be mediated by changes in soil-related parameters (Fig. 1, *paths 3* and *8*), for example, soil compaction [12,15], which results in bare soil exposure and restricts the ability of roots to penetrate and of shoots to emerge [12], therefore preventing the restoration of vegetation [16]. Furthermore, soil compaction may affect the availability of water and nutrients to plants [15,17] (Fig. 1, *paths 5* and *path 8*) through the altered decomposition processes via reduced soil porosity [15] (Fig. 1, *path 10*) or decreased activity of soil decomposers [18] (Fig. 1, *path 6*), which in turn are known to impact nutrient cycling and, thereby, resource availability for plants [19] (Fig. 1, *paths 5, 7* and *8*). On the other hand, the deposition of excrements by cattle in soil can relax soil acidification [12] (Fig. 1, *path 9*), and, as a result, can increase the nutrient uptake by plants [15,20] (Fig. 1, *paths 8*). Furthermore, cattle manure serves as the resource input for soil decomposers [17] (Fig. 1, *path 4*), therefore resulting in increased soil organic matter content [21] (Fig. 1, *path 7*), and ultimately in greater resource availability to plants (Fig. 1, *paths 8* and *5*). Moderate levels of soil disturbance by trampling can also increase water and nutrient availability in vegetation gaps [13] and may stimulate germination of plants from the soil seed bank as a result of increased light availability and nutrient-rich and pathogen-free soil [14] (Fig. 1, *path 3*).

Grazing effects on vegetation patterns may vary strongly with elevation and topography. Livestock density is usually lower at higher elevation (Fig. 1, *path 12*) leading to lower net effects of grazing on the grassland community. On other hand, an extensive erosion processes in steep slopes of terrain with heavy runoff and soil losses limit water and nutrient availability to plant growth [22]. Thus, the use of vegetation by livestock is expected to have more pronounced effects on plant diversity and composition in uplands in contrast to those in lowlands [23] (Fig. 1, *path 15*). Furthermore, soil types vary with topography (Fig. S1). This might alter soil properties (i.e., pH and soil organic carbon) along the elevation gradient [24] (Fig. 1, *path 13*), therefore influencing nutrient availability to plants (Fig. 1, *paths 8, 15* and *5*). In addition, higher soil leaching with increasing elevation is also expected to increase soil acidity and decrease soil organic carbon [24]. On the other hand, increased elevation involves altered climate conditions (i.e., increased solar radiation, precipitation, humidity, extensive wind exposure, and reduced air temperature), which may further affect local vegetation by shaping the regional species pool composition, i.e. via filtering of species physiologically capable of living under these environmental conditions [25] (Fig. 1, *path 11*). High topographic variability linked to steep altitudinal gradients across mountain areas are found to result in greater habitat diversity [26], and therefore may lead to increased plant species richness along the elevation gradient (Fig. 1, *path 11*). Increased precipitation with elevation in the Carpathians [27] may also alter the effects of grazing on plant diversity [28].

The variety and interacting effects of these different mechanisms may be responsible for the idiosyncratic nature of grazing effects on plant communities in grasslands. Given that a single mechanism may produce multiple patterns (Table S1), while multiple mechanisms may lead to convergent patterns or tradeoffs among different effects, it has been shown that the relationships among grazing and biodiversity are best understood within the context of multivariate models [29]. In fact, recent studies have started to reveal the variety of pathways via which grazing affects plant community in grasslands [10,30–32]. However, the majority of the existing evidence is limited to contrasting grazed *vs* ungrazed (i.e., fenced) systems. Studies on the effects of realistic grazing situations, where grazing managements varies from low to high intensities, are largely under-represented. This limits our ability to predict grazing effects on natural or semi-natural grasslands where fencing is uncommon.

In this study, we evaluate the simultaneous direct and indirect effects of grazing intensity and elevation on plant species richness, number of key plant functional groups (legumes, grasses (Poaceae), other monocots (rushes and sedges), and forbs), and on the proportion of undesirable weeds in semi-natural grasslands at large topographic scale ranging from the Carpathian Mountains in Ukraine, across the adjacent foothills to the plain areas (Fig. 2), which is a largely under-represented region in scientific literature. We test concomitantly whether the effects of grazing intensity and elevation on plant community operate via changes in soil properties and altered soil biota; and if grazing impact on plant diversity varies across the elevation gradient (i.e., an interactive effect of grazing and elevation, Fig. 1, *path 16*).

**Fig. 2.**
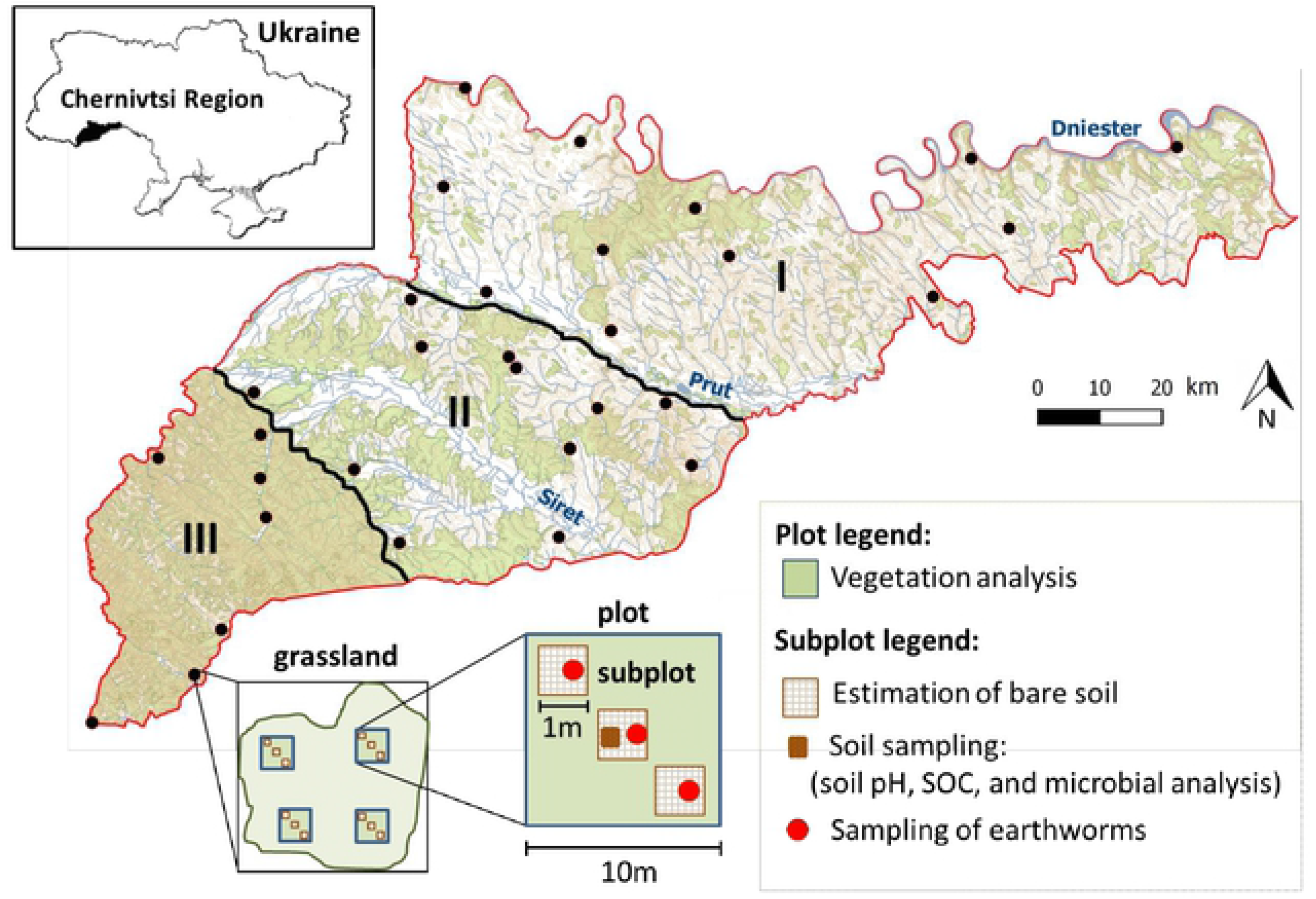
Location of studied grassland sites (filled dots, n=31) across the study area (Eastern Carpathians, Chernivtsi Region, Ukraine) and experimental design. Physical-geographical zones: I plains (n=12 grassland sites); II foothills (the Precarpathians) (n=12 grassland sites); and III mountains (n=7 grassland sites). Green areas show forest habitats; and white areas show unforested lands. The rivers Dniester, Prut, and Siret are major rivers of Chernivtsi Region. SOC, soil organic carbon. The map was created using QGIS 2.18.26 (https://qgis.org). The forest layer is based on the Public cadastral map of Ukraine (https://map.land.gov.ua). Four plots (10 m × 10 m) per grassland site and three subplots (1 m × 1 m) within each plot were selected.

## Material and Methods

### Study area

We studied 31 semi-natural grasslands (in 2006 and 2007), that have been used for decades as public pastures for cattle grazing. The study grasslands are distributed throughout the Chernivtsi Region (47°43’ − 48°41’ N × 24°55’ − 27°30’ E) located in the south-west of Ukraine along the rivers Dniester, Prut and Siret (Fig. 2). Pastures account approximately for 15% of the agricultural land area in the region. The study area generally experiences a temperate humid continental climate, with a significant amount of precipitation, which increases with elevation (see below for details). Although the Chernivtsi Region is the smallest within Ukraine by land area (8,097 km^2^), it is highly diverse in environmental conditions. According to the specific characteristics of landscapes, climate, elevation, and age and type of rocks it can be divided into the three physical-geographical zones: *the plains*, *the foothills* (the Precarpathians) and the *mountains* (the Carpathian Mountains) (Fig. 2) [33].

The plains of the Chernivtsi Region are located within the southwestern edge of the East European Plain. The landscape is mainly flat and densely dissected by river valleys and ravines. The plains represent ~49% of the territory of the region (Fig. 2) and are located between the rivers Prut and Dniester (the Prut-Dniester interflow). The average elevation is 230 m a.s.l. with the highest elevation 515 m a.s.l. on hill formation in the center of the zone. The average annual rainfall is 575–660 mm, and the mean annual temperature is 8 °C with the averages of −5 °C in January and 20 °C in July. The soils are mostly Luvisols (pH=5.2−6.5, humus=1.5−4%), Phaeozems (pH=5.5−6.8, humus=4−6%), and Chernozems (pH=6.5−7, humus=5−12%) (Fig. S1). A high level of summer precipitation causes frequent soil erosions. The plains zone is characterized by deciduous forest vegetation (*Fagus sylvatica*, *Quercus robur*, and *Carpinus betulus*) [33]. In the west of the area there are meadow steppes with *Festuca valesiaca*, *Poa angustifolia*, *Carex humilis*, and *Brachipodium pinnatum*, each dominating in different patches [33].

The Precarpathians (the foothills) cover ~26% of the Chernivtsi Region and are located in the center of the region (Fig. 2). The Precarpathians are formed as a part of the foredeep at the outer base of the Carpathian Mountains (i.e., part of the Outer Carpathian Depression). They are considered the foothills of the Ukrainian Carpathians and represent a transition from the plains to the mountains. The terrain is highly complex due to the predominance of the wavy hills, densely dissected by the rivers Prut and Siret. The average elevation is 350 m a.s.l. (rising up to 537 m a.s.l.); and the average annual rainfall is 575-780 mm. The mean annual temperature is 7 °C with the average temperatures in January −4.5°C and in July 18 °C. The soils are mostly Retisols (pH=4.3−4.5, humus=1−3%) (Fig. S1). Luvisols soils also can be found over the entire foothills (Fig. S1). The vegetation mostly consists of broad-leaved forests (*Fagus sylvatica*, *Carpinus betulus*, *Acer pseudoplatanus*), mixed fir-spruce (*Abies-Picea*) and fir-beech (*Abies-Fagus*) forests, and mixed forb − graminoid meadows [33].

The mountain zone covers ~25% of the Chernivtsi Region and is part of the Carpathian Mountains (i.e., the Outer Eastern Carpathians) with an average elevation of about 900 m a.s.l. (from ~600 to 1565 m asl). The climate is colder (mean annual temperature: 4.6 °C with the average temperature in January −7.4 °C and in July 15 °C) and excessively humid (annual rainfall: 700−1200 mm) in comparison with the other zones. The relief is a typical mountain relief with a variety of landscapes such as low-hill terrains covered by forests and secondary meadows; mountains with a medium elevation covered by forests; and subalpine mountains covered by subalpine meadows. Soils are mostly Cambisols (рН=4.3−5.5, humus=1−8%), and Retisols (Fig. S1). The prevailing vegetation closely follows the elevation lines: European beech (*Fagus sylvatica*) and beech-fir (*Fagus-Abies*) forests at 800–950 m asl; fir-spruce (*Abies-Picea*) forests at 950–1100 m asl; spruce (*Picea abies*) forests at 1100–1400 m asl; and subalpine meadows “polonynas” and shrub-lands > 1400 m a.s.l. [33].

### Site selection

Of the 31 grasslands sites, 12 were selected within the plains, 12 within the foothills and 7 within the mountains. All grasslands have been used as common grazing lands for cattle pasturing by private households, which typically have 2-3 livestock units per household (for calculations of livestock density in units per area see explanations below and Table S2). Grazing season depends on the growing season of vegetation and varies for the three physical-geographical zones. The grazing season for mountain grasslands lasts nearly 120-150 days, for the foothills around 180 days and for the plains zone 210-220 days.

Sampling was performed identically for each of the compared ecosystems during June–July in 2006 and 2007. The same grassland sites were sampled twice, i.e. one time per year. A handheld GPS-12 Garmin^®^ was used to identify the geographic coordinates and average elevation (m a.s.l.) for each study ecosystem. Four plots (10 m × 10 m) were selected within each of the 31 grassland sites (Fig. 2). Placement of plots was random but constrained by the edges and size of the field site. The average closest distance between two neighboring plots was 10 m and was chosen to minimize the potential for spatial autocorrelation influencing the results. The distance from plot to the edges of the grassland was 10 m to prevent edge effects. Within each plot, a transect was positioned diagonally through the plot. Three 1 m by 1 m subplots for estimation of bare soil, and earthworm sampling were randomly selected along the transect with a minimum distance to the next subplot of 1 m (Fig. 2). Soil samples for microbial and chemical analysis were taken from one of the three subplots within each plot (Fig. 2). Locations of plots and subplots within each grassland site differed across the two years.

### Cattle density and bare soil exposure

Cattle density was measured as the number of livestock units per hectare of grassland area (livestock units h^−1^) (Table S2). For this, the number, type and age of cattle were recorded for private households, which used the pastures during the grazing seasons of the two study years. We transformed the number of cattle to livestock units based on the widely used conversion factors for Europe (Table S2) and used this cattle density measure as a proxy for the degree of grazing intensity of the study pastures. For each 1 m² subplot, the fraction of bare soil was visually estimated. The mean of the three subplots and then the four plots was taken to approximate the percent bare soil per grassland in each year. Cattle density and the fraction of bare soil were averaged across the two sampling years.

### Vegetation

Vegetation was recorded within each plot during the peak growing season for the different physical-geographical zones (June–July) in the two study years. All plant species were determined within each plot. We used the number of species as a measure of plant species richness 100 m^−2^. Within each plot three 1 m × 1 m randomly selected subplots were used to estimate the relative cover of each species by vertically projecting canopy cover (%) for each species within each subplot. Averages were then taken across the three replicate subplots. All recorded species were classified into four functional groups: legumes (Fabaceae), grasses (Poaceae), other monocots (rushes and sedges), and forbs (other than legumes). The number of functional groups was used as the measure of plant functional diversity per 100 m^2^. Further, species were classified as undesirable weeds (Table S4) if they were known to reduce grazing efficiency, forage yield, palatability and quality, therefore contributing to lower forage and animal production of grassland ecosystems [34]. The group of undesirable weeds includes both unpalatable species as well as competitive weeds. Unpalatable plants are those containing toxic compounds poisonous to cattle (e.g., *Equisetum arvense*, *Ranunculus acris*, *Saponaria officinalis*, *Euphorbia* sp.), or, when eaten, may cause mechanical injuries because of a spiny covering or fine hairs (e.g., *Carduus crispus*). Competitive weeds (e.g., some coarse tall grasses and forbs) are not toxic to cattle and somewhat palatable (e.g., *Plantago* sp.), but can increase in density over time and outcompete desirable forage species. In a pasture, they reduce grazing efficiency of cattle by increasing search time for food. The number of undesirable weed species was quantified on the 100 m^2^ plots. Data on plant species richness, functional diversity and richness of undesirable weeds were averaged across the four plots level for each year and further averaged across the two sampling years. Further, we measured the proportion of undesirable weeds as the ratio of the species number of the undesirable weeds to the total plant species number. Data on canopy cover (%) for each species were averaged across the four plots within each year and scaled to unitless relative cover measures (C) ranging from 1 to 5 in accordance with Braun-Blanquet [35]: C = 1 for cover from 1 to 5 %; C = 2 for cover from >5 to 25 %; C = 3 for cover from >25 to 50 %; C = 4 for cover from >50 to 75 %; C = 5 for cover >75%. Further, the relative cover measures for each species were averaged across the two sampling years.

### Soil sampling and analysis

Before soil sampling, vegetation and upper litter layer were removed in a small area. One soil sample per plot (for a total of four samples per grassland, Fig. 2) was collected at 0−10 cm depth during the vegetation sampling campaign within each of the grassland sites using a soil corer with a diameter of 5 cm. The soil samples were immediately stored at −5 °C for microbial and chemical analysis. For the examination of organic carbon content and soil pH analysis the soil samples were air-dried, sieved (mesh width 2 mm) and homogenized. Soil organic carbon (%) was determined using a Tyurin’s wet combustion technique, which is based on organic carbon oxidation by potassium dichromate (0.4 N) in acid solution (K_2_Cr_2_O_4_: H_2_SO_4_ in a 1:1 ratio). The soil pH was determined by a standard glass electrode pH meter using a potassium chloride solution in a 1:2 ratio (soil : 0.1-N KCl).

### Density of soil biota

For the soil microbiological analysis, we counted cells of three microbial groups: heterotrophic bacteria, micromycetes, and actinomycetes. Cells were cultured on group-specific substrates under controlled temperature conditions: heterotrophic bacteria were cultured on meat-peptone agar between 28 and 30 °C, micromycetes were cultured on modified Czapek-Dox substrate with streptomycin at 20 to 25 °C, and actinomycetes were cultured on starch-ammonium agar at 28 to 30 °C. The total number of cells of all three groups was used as abundance measure for the soil microbial community (cells × 10^8^ g^−1^ dry soil). We took averages across the four plots (Fig. 2) to approximate the abundance per grassland for each year. Further, the abundance data were averaged across the two sampling years. To sample earthworms we used a standard Quantitative Hand Sorting method. For this, 30×30 cm^2^ soil blocks with a depth of 15 cm were excavated from each of three subplots in each of four plots (leading to 12 samples per grassland, Fig. 2), and earthworms were immediately separated manually in the field and sampled into empty vials. At the same day specimens were counted in the lab, their fresh weight was determined, then they were oven-dried and their dry weight was determined. Earthworm dry weight data were calibrated to an area of 1 m^2^. Averages were taken across the four plots to calculate earthworm biomass per grassland (g m^−2^) for each year and further averaged across the two sampling years.

### Data analysis

All analyses were performed in R version 3.4.3 [36]. We applied structural equation modelling (SEM) techniques using the package ‘lavaan’ [37] in R as an exploratory approach for assessing direct and indirect simultaneous effects of grazing intensity on plant species richness, functional diversity and proportion of undesirable weeds along the elevation gradient (Fig. 1). Plant species richness and plant functional diversity were log-transformed; and cattle density was square-root transformed to meet normality of errors and homoscedasticity. We used a Chi-square (*χ*^2^) test with maximum likelihood ratio, Root Mean Square Error (RMSEA), Comparative Fit Index (CFI) and Tucker–Lewis Non-Normed Fit Index (NNFI) and Standardized Root Mean Square Residual (SRMR) as goodness of fit tests to assess the validity of the SEM model [38]. The individual effects included in the model were evaluated for significance (the *P*-value is lower than α = 0.05), and standardized SEM regression coefficients were used as a quantitative measure of the strength of these effects (Table S6, Fig. 2). We also tested the significance of the mediation effects, i.e. all indirect paths (effects among two variables mediated by other variables) were also evaluated for significance (Table S7). To assess whether the grazing-intensity effects on plant community varied across the elevation gradient, we tested for an interactive effect of grazing and elevation (Fig. 1, *path 16*) on each of the response variable, i.e. on plant species richness, functional diversity, and the proportion of undesirable weeds. The interactive effects of grazing and elevation (Fig. 1, *path 16*) were not deemed significant in the original SEM model (Fig. S2), and they were removed in the final model (Fig. 3) to achieve adequate fit statistics. The final SEM model (Fig. 3, Table S6) was well supported by the data (χ^2^ = 10.2, df = 9, P = 0.33, RMSEA = 0.07, P_RMSEA_ = 0.39; CFI = 0.99; NNFI = 0.96; SRMR = 0.05), and an addition of any other paths did not improve the model, suggesting that all important relationships were specified To investigate plant community composition we used nonmetric multidimensional scaling (NMDS) based on Bray–Curtis matrices using the ‘metaMDS’ function in the *vegan* package in R with a maximum of 100 random starts. We plotted differences in community composition using an NMDS plot and fitted grazing intensity and elevation post hoc using the ‘envfit’ function and the ‘ordisurf’ function, respectively. To test the effects of elevation, cattle density, soil properties, and soil biota on plant community composition we performed a PERMANOVA test on Bray-Curtis matrices with 1000 permutations using the ‘adonis’ function in the *vegan* package in R.

**Fig. 3.**
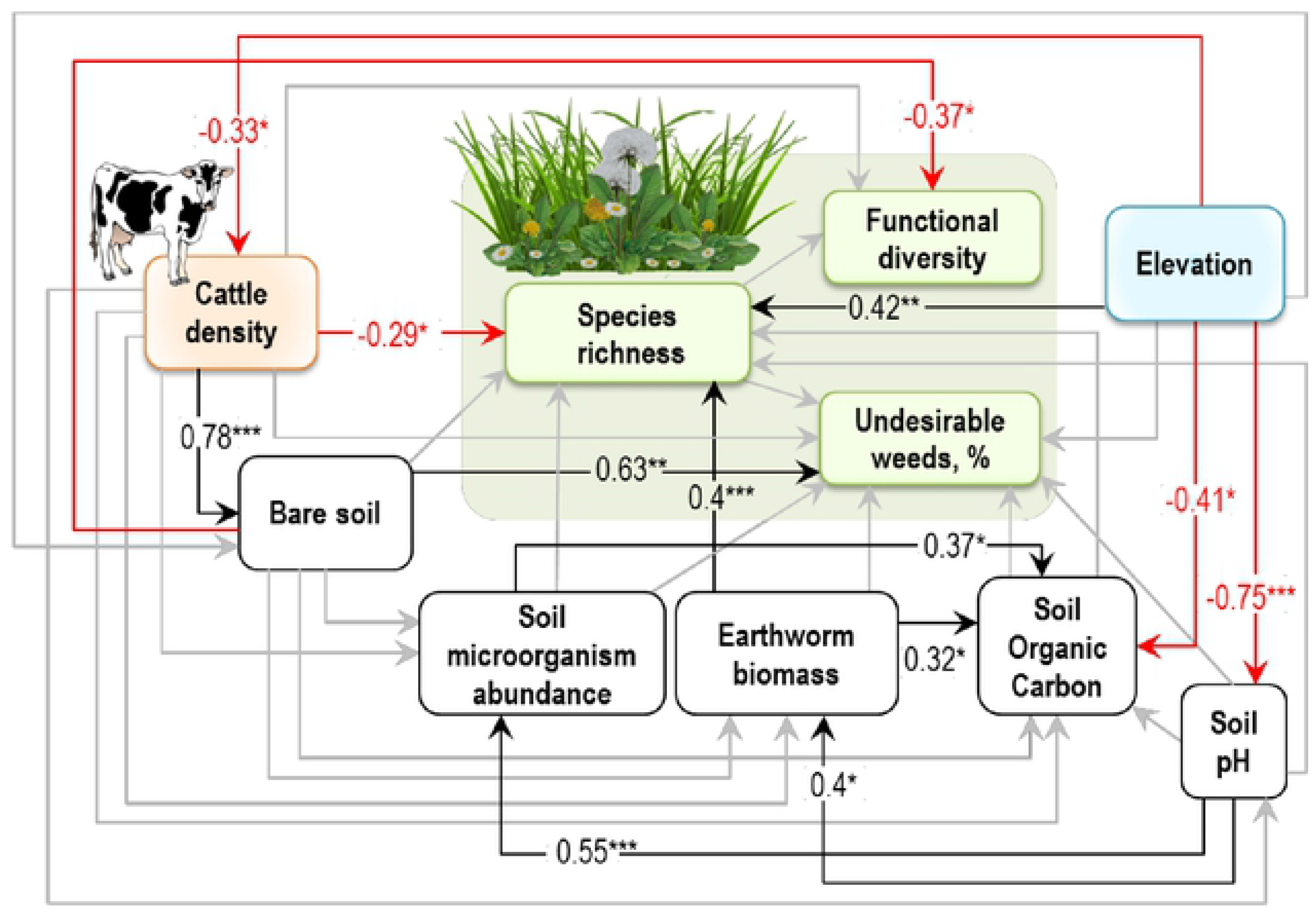
SEM showing the direct and indirect pathways through which grazing intensity and environmental conditions affected plant diversity and composition. Models were well supported by our data (χ^2^ = 10.2, df = 9, P = 0.33, RMSEA = 0.07, P_RMSEA_ = 0.39; CFI = 0.99; NNFI = 0.96; SRMR = 0.05). Numbers associated with each arrow are standardized path coefficients with the following levels of significance: *P ≤ 0.05; **P ≤ 0.01; ***P ≤ 0.001. Red and black paths represent significant (i.e., P ≤ 0.05) negative and positive path coefficients, respectively, and grey paths were not significant. RMSEA: Root Mean Square Error of Approximation; CFI, Comparative Fit Index; NNFI: Tucker–Lewis Non-Normed Fit Index; SRMR: Standardized Root Mean Square Residual. See Tables S6 for more details.

## Results

In the 31 grasslands (Fig. 2) we sampled a total of 175 plant species: 13 legumes (Fabaceae), 18 grasses (Poaceae), 8 other monocots (i.e., rushes and sedges), and 136 non-legume forbs. Plant species richness varied from 7 to 51 species per grassland (Table S3). The list of species and summary statistics of the richness derived from the plant community data for each of the physical-geographical zones are given in Table S3-S4. In total, we classified 86 species as undesirable weeds (thereof 48 species in the plains, 60 species in the foothills, and 62 species in the mountains, see Table S3), which varied from 5 to 28.5 species per grassland (13.9±1.08; mean±s.e.m.). Cattle density of the study grasslands varied from low (0.01 livestock units h^−1^) to high (1.38 livestock units h^−1^), and elevation varied from 399 to 1323 m a.s.l. across the grassland sites (Table S5).

### Drivers of plant species richness

Overall, the sampled plant communities were rich in species number (27.5±1.97, mean±s.e.m.), reaching a maximum of 51 species per grassland site (Table S3). We found that cattle density significantly reduced plant species richness (Fig. 3). Disentangling direct and indirect effects using SEM (Table S7) revealed only direct impacts of cattle density on species richness, while none of the indirect effects (i.e., mediated via bare soil exposure, soil properties, or soil decomposers) were significant (Fig. 3, Table S7). In contrast, elevation directly and indirectly influenced plant species richness in our grassland sites. The direct effect was strongly positive while the indirect effect was negative, resulting in a weaker positive overall effect. The indirect negative effect of elevation was mediated via soil pH and earthworm biomass (Table S7). Specifically, a decrease in soil pH with increasing elevation reduced earthworm biomass, which in turn resulted in lower plant species richness (Table S7).

Cattle density decreased with elevation in our grassland sites (Fig. 3), however, grazing-mediated indirect effects of elevation on plant species richness were not significant (Table S7). Furthermore, effects of cattle density on species richness did not vary across the elevation gradient, as the interactive effects of grazing and elevation on plant species richness were not significant (Fig. S2). In general, at our grassland sites the overall impact (direct and indirect effects) of cattle density on plant species richness was stronger than that of elevation (Table S7).

### Drivers of functional diversity and community composition

All significant effects of cattle density on plant functional diversity and on percent of undesirable weeds in plant community were indirect and mediated by bare soil exposure (Table S7). Specifically, greater cattle density led to an increased fraction of bare soil, which in turn decreased plant functional diversity (Fig. 3). In contrast, grazing-induced bare soil increased percentage of undesirable weeds (Fig. 3), therefore resulting in positive overall impact of grazing intensity on the proportion of undesirable species in a plant community. Cattle density reduced species number of legumes. Species number of non-legume forbs and of rushes and sedges increased with increased fraction of bare soil under greater grazing intensity (Table S8). In contrast, species number of grasses was not influenced by cattle density in our grassland sites.

The SEM model indicated no significant overall effects of elevation on functional diversity or on the proportion of undesirable weeds in a plant community (Table S7). When testing the effect on each functional group separately in linear models, we found a positive effect of elevation on the species number of rushes and sedges (Table S8). There were no significant effects of elevation on species number of legumes, non-legume forbs, or grasses (Table S8). The interactive effects of grazing and elevation on plant functional diversity and on proportion of undesirable weeds were not significant (Fig. S2).

We found no significant effects of species richness on functional diversity and on the proportion of undesirable weeds (Fig. 3). However, species number of legumes increased with both, community species richness and functional diversity. Species numbers of both grasses and other monocots (i.e., rushes and sedges) were significantly higher in more functionally diverse plant communities but not affected by community species richness. In contrast, species number of non-legume forbs was positively affected by community species richness but negatively affected by functional diversity (Table S8).

Plant community composition was assessed based on individual species’ canopy cover. The changes in community composition are shown in the NMDS plot (Fig. 4). Both changes in elevation and grazing intensity significantly affected plant community composition (elevation: F_1,30_ = 1.96, R^2^ = 0.06, P = 0.005; grazing: F_1,30_ = 1.63, R^2^ = 0.05, P = 0.03), while the interactive effects of grazing and elevation and soil parameters were not significant (Table S9).

**Fig. 4.**
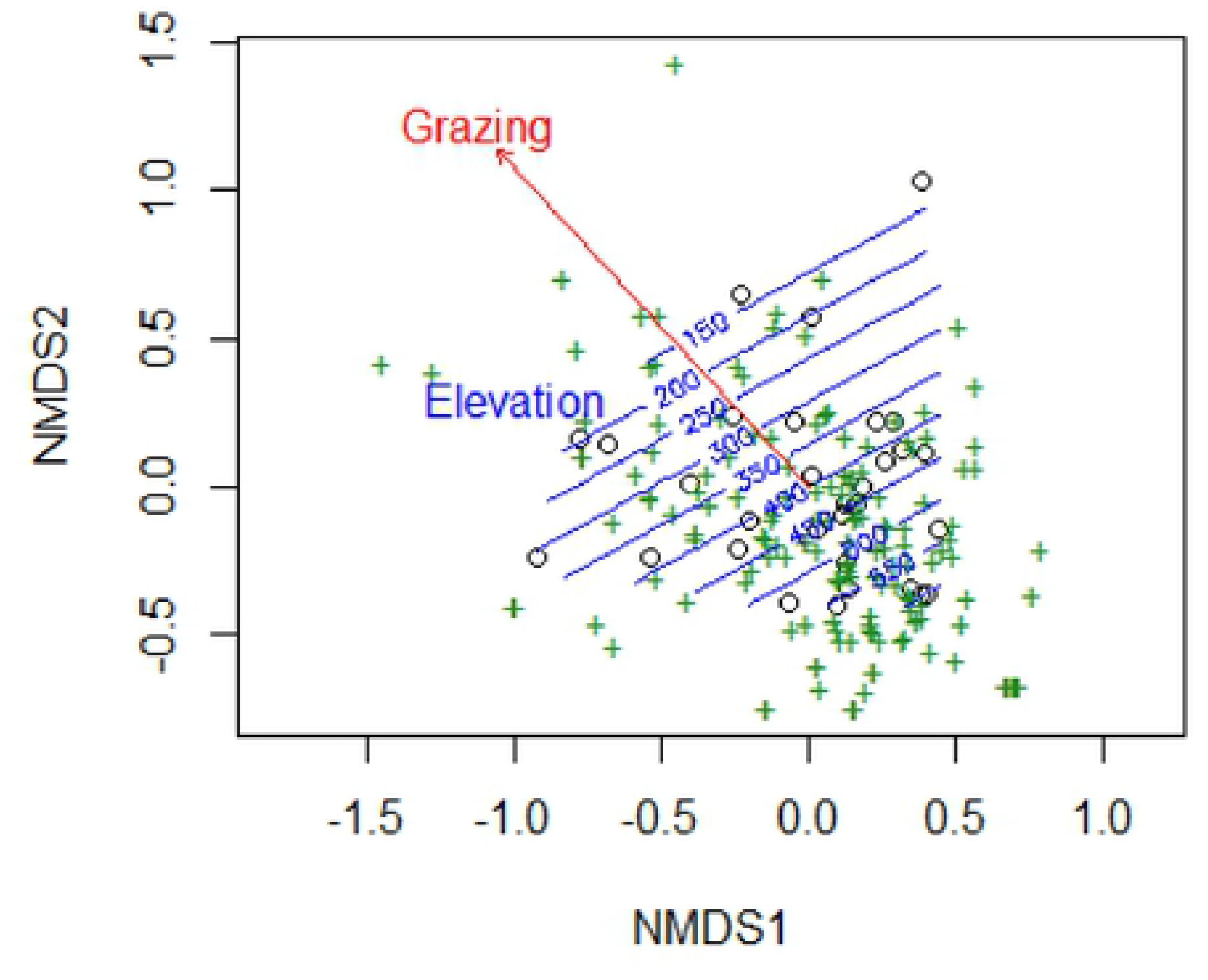
Nonmetric multidimensional scaling (NMDS) plot showing the composition of plant communities in grasslands. Symbols in green show species (n = 175), symbols in black show grasslands (n = 31). Grazing intensity and elevation significantly affected community composition (see Table S9). Grazing intensity is shown by a red arrow drawn by fitting cattle density post hoc. Elevation gradient is illustrated by the blue contour lines. Stress = 0.16.

## Discussion

Our investigation of plant diversity drivers in temperate semi-natural grasslands at a relatively large topographic scale showed strong negative effects of cattle density on plant species and functional diversity accompanied with positive effects on the proportion of undesirable weeds. Structural equation modelling revealed that one of these effects was directly linked to grazing (i.e., species richness), while others (i.e., functional diversity and fraction of undesirable weeds) were mediated by bare soil exposure. Elevation had a strong positive direct effect on plant species richness and a negative indirect effect mediated via altered soil pH and earthworm density. Both elevation and grazing shifted plant community composition based on individual species’ canopy cover. Although grazing intensity decreased with elevation, the effects of grazing on plant diversity or community composition did not vary across the elevation gradient.

### Effects of grazing

In our study grazing affected plant species richness, functional diversity, the proportion of undesirable weeds in the plant community (Table S7) and community composition (Table S9). However, the underlying mechanisms of these effects differed: Increasing grazing intensity reduced plant species richness and this effect was driven mainly by a direct link between cattle density and plant species richness (Fig. 3), while none of indirect effects here were significant (Table S7). These findings are in agreement with Socher et al. [39] who also found that effects of grazing on plant species richness were governed by direct effects, in contrast to other land-use types, e.g. fertilization. This direct effect can likely be explained by the mechanical destruction of vegetation, overgrazing, and chemical and biological impact of faeces and urine depositions [12] (Table S1), but possibly also by an altered colonization of species from regional species pools via the removal of seeds and reproductive structures of plants by grazing [40].

In contrast to species richness, the effects of grazing on functional diversity and on the proportion of undesirable weeds were completely mediated via bare soil exposure (Fig. 3). Specifically, we found a reduction in functional diversity and an increase in the proportion of undesirable weeds along the increased fraction of the grazing-induced bare soil (Fig. 3, Table S7). The observed reduction in functional diversity was a result of the loss of legume species from the plant community under increasing grazing intensity (Table S8). Most of legume species observed in our study are palatable to livestock (Table S4), and may perhaps be favored among the other plant groups for grazing. Previous studies show that palatable legumes have generally lower persistence under grazing in contrast to other functional groups [41], due to the preferential grazing of forage legumes by ruminants [42]. Furthermore, high inputs of nitrogen in the soil, delivered with livestock urine [12], have been previously found to suppress legumes [43]. Instead, the less palatable and more competitive species might further benefit from the grazing pressure, which may create favorable conditions for their seed germination [13,44]. Our results are aligned with this prediction as we found that competitive undesirable weed species, which are predominantly rushes, sedges and some non-legume forbs (Table S4), increased in species number with increased fraction of cattle-induced bare soil (Table S8). This observed pattern confirms findings of other studies. For example, Tardella [45] found that the presence of bare soil, linked to lower soil depth, increased the number of stress-tolerant and ruderal species in a plant community. Similarly, Symstad [46] showed that an increased fraction of bare soil promotes the invasion success of highly competitive species. This general finding of a shift in plant community composition towards undesirable weed species under grazing can be attributed to several possible mechanisms. Compacted by cattle trampling bare soil reduces nutrient and water availability to plants [15,20] and restricts the ability of plant seedlings to penetrate the soil and to emerge [12]. This may lead to a shift in community composition by favoring species with high competitive abilities for resource acquisition. On the other hand, an increased proportion of bare soil with increasing cattle density might lead to more pronounced spatial heterogeneity due to increased vegetation patchiness [14]. Such open canopy gaps then filter for adapted colonizer species [47], for example short-lived forbs [46,47]. Thus, the tendency of undesirable weed species to increase in grazing-induced bare soil patches in our study may relate to both, their competitive ability for resources and to their ability of colonizing bare soil [47].

### Effects of elevation

We found a strong effect of elevation on plant community composition (Table S9) and diversity in our grassland sites (Fig. 3, Table S7). Conversely, we found no significant effects of elevation on functional diversity of the plant community or on the proportion of undesirable weeds (Table S7). An observed positive effect of elevation on the species number of rushes and sedges (Table S8) might be attributed to the increased humidity with elevation [24], as rushes and sedges are known to dominate other species in increased soil moisture conditions [48]. Elevation was found to impact plant diversity in previous studies [29,49,50], mainly because of the correlation with climatic factors [29,50]. Increased precipitation along the elevation gradient in the Carpathians [27,51] is associated with increased site productivity [10], which in turn has been shown to increase plant diversity [28]. Additionally, higher topographic variability with increasing elevation across our study area results in greater habitat diversity [26] and thus may allow plants to find suitable habitats within small distances, therefore potentially leading to higher species richness.. On the other hand, our results also show that the effect of elevation on plant species richness was partially negative and mediated by reduced biomass of earthworms as a result of increased soil acidification with elevation (Fig. 3, Table S7). The observed decrease in soil pH with elevation in our study could be linked to higher soil leaching processes in more humid and lower temperature conditions with increasing elevation [24]. Furthermore, within our study area, the change in climatic conditions with elevation and topography determines the differences in the soil types, which further result in variation of soil properties along the elevation gradient, including soil pH [24]. Increased soil acidity was found to suppress decomposition, burrowing, casting and mixing activities of earthworms with cascading consequences for growth [18] and competitive interactions among plant species [19].

Elevation had a weak impact on cattle density in our grassland site (Fig. 3), leading to nonsignificant mediation of elevation effects on plant species richness, functional diversity, and on proportion of undesirable weeds through grazing (Table S7). Furthermore, we found that the interactive effects of grazing and elevation on plant diversity and composition (Fig. S2) were not significant. These results indicate that the effects of grazing intensity on plant community in our grassland sites did not change across the elevation gradient. Compared to grazing, elevation expressed a relatively stronger direct effect on plant species richness in our grassland sites (Fig. 3). This is in agreement for example with Grace and Pugesek [49] and Báldi et al. [6] reporting that abiotic conditions exerted a relatively strong control over species richness, with grazing playing a less important role. However, the opposing direct and indirect effects of elevation on plant species richness in our grassland sites led to weaker positive overall effect. Furthermore, the same signs of the direct and indirect effects of grazing on plant species richness led to strong negative overall effect. As a result, an overall impact of cattle density on plant species richness was stronger than that of elevation in our grassland sites (Table S7).

### Consequences for ecosystem resilience, functions and services

The variation in plant diversity and composition caused by grazing along the environmental gradients reported here may have consequences for ecosystem functioning and services mediated by land use [3,9,52]. Cattle production itself is a valuable service but if other ecosystem services of the grasslands are to be considered, cattle grazing affect them. Our results show that high livestock stocking rates can directly induce changes in plant species richness, therefore likely influencing the availability and heterogeneity of resources to a wide range of organisms with possible cascading effects on biodiversity across trophic levels [9,53,54] and related ecosystem processes [52], such as primary productivity [32], herbivory, decomposition, and predation [19,55,56]. Furthermore, the physical impact of the heavy livestock on soil resulted in bare soil exposure in our grassland sites with some of the most consistent outcomes for plant community composition being reduced plant functional diversity and increased proportion of undesirable weeds. This is likely to reduce grazing efficiency, forage yield and quality, therefore contributing to lower forage and animal production of grassland ecosystems [34]. Our results on a grazing-induced increase of bare soil entail important implications for the management of ecosystem functions and service, since previous studies show that bare soil exposure increases soil erodibility [22], leaching and plant invasion processes [46,57], loss of nitrogen and pollution of the atmosphere with N_2_O during denitrification of waterlogged areas [12] and alters ecosystem watershed function [5].

Most of the Carpathian’s unique biodiversity is dependent on semi-natural grasslands [58], which in turn require regular grass removal, such as via grazing, in order to survive [2]. However, as our results indicate, the benefits of grazing depend on the grazing intensity. As shown by our data, while low levels of grazing create high-diversity plant communities, high stocking rates reduce plant species and functional diversity and alter plant composition. On the other hand, farmland abandonments all over Europe, ongoing for least three decades [58] lead to the replacement of grasslands with successional shrublands, thus reducing ecosystem biodiversity and functioning [3]. Furthermore, our results indicate that grazing effects on soil and related ecosystem functions and services in semi-natural grasslands are likely more pronounced along the elevation gradient. The reduced soil organic carbon with elevation shown by our data (Fig. 3) might deteriorate the soil’s ability to resist trampling pressure by cattle in our grassland sites [12] and to decrease grassland productivity [32]. We also found a lower density of earthworms and soil microorganisms with elevation, potentially impacting nutrient cycling, resource availability for plants, and plant productivity of grasslands [18].

## Conclusions

The current study provides data from a largely under-represented region in the scientific literature, the Eastern Carpathians in Ukraine. If we extrapolate our results to the entire region, they suggest implications for land-use management strategies of the Carpathian semi-natural grasslands, where preserving biodiversity is crucial for the maintenance of ecosystem functioning and provisioning of ecosystem services [5–7,52]. Our main findings are that plant diversity and composition are controlled by the complex interplay among grazing intensity and environmental conditions along the elevation gradient. Both, grazing and elevation affected plant community directly and indirectly via altered soil properties and soil biota. Thus, the assessment of land-use effects on grassland biodiversity should encompass simultaneous observations of multivariate parameters and links among them. Maintaining low levels of grazing in the semi-natural grasslands and taking into account the ecosystem characteristics driven by elevation and topography are therefore essential for maintaining species-rich and functionally diverse plant communities.

## Acknowledgements

We are grateful to Vasyl Budzhak (Department of Botany, Forestry and Landscape Architecture, Chernivtsi National University) for help with species identification, to Christoph Scherber (University of Muenster) for discussions on the conceptual model, and to all colleagues from the Department of Ecology and Biomonitoring (Chernivtsi National University) and Theoretical Ecology Group (Freie Universität Berlin) for discussions. We thank Stepan S. Kostyshyn for management support and for facilities provided for field work and laboratory analyses. Further, we acknowledge the technicians and student helpers for their work.

## Author contributions

OYB and JSP conceived and designed the study. OYB and RSS contributed data. OYB analyzed the data. VAN designed the maps. OYB wrote the original draft, and BT, JSP, RSS and VAN substantially contributed to discussions, review and editing.

## References

1. Food and Agriculture Organization of the United Nations. Grasslands of the World. Plant Production and Protection Series. [Internet]. 34th ed. Suttie JM, Reynolds SG, Batello C, editors. Rome: Food and Agriculture Organization of United Nations; 2005. Available: http://www.fao.org/docrep/008/y8344e/y8344e00.htm#Contents

2. WallisDeVries MF, Poschlod P, Willems JH. Challenges for the conservation of calcareous grasslands in northwestern Europe: Integrating the requirements of flora and fauna. Biological Conservation. 2002;104: 265–273. doi:10.1016/S0006-3207(01)00191-4

3. Soliveres S, van der Plas F, Manning P, Prati D, Gossner MM, Renner SC, et al. Biodiversity at multiple trophic levels is needed for ecosystem multifunctionality. Nature. Nature Publishing Group; 2016;536: 456–459. doi:10.1038/nature19092

4. World Resources Institute. A Guide to World Resources 2000-2001: People and Ecosystems: The Fraying Web of Life. World Resources Institute; 2000.

5. De Keersmaecker W, van Rooijen N, Lhermitte S, Tits L, Schaminée J, Coppin P, et al. Species-rich semi-natural grasslands have a higher resistance but a lower resilience than intensively managed agricultural grasslands in response to climate anomalies. Journal of Applied Ecology. 2016;53: 430–439. doi:10.1111/1365-2664.12595

6. Báldi A, Batáry P, Kleijn D. Effects of grazing and biogeographic regions on grassland biodiversity in Hungary - analysing assemblages of 1200 species. Agriculture, Ecosystems and Environment. Elsevier B.V.; 2013;166: 28–34. doi:10.1016/j.agee.2012.03.005

7. Turtureanu PD, Palpurina S, Becker T, Dolnik C, Ruprecht E, Sutcliffe LME, et al. Scale- and taxon-dependent biodiversity patterns of dry grassland vegetation in Transylvania. Agriculture, Ecosystems and Environment. 2014;182: 15–24. doi:10.1016/j.agee.2013.10.028

8. Sala OE, Chapin III FS, Armesto JJ, Berlow E, BloomÞeld J, Dirzo R, et al. Global biodiversity scenarios for the year 2100. Science. 2005;287: 1770–1774. doi:10.1126/science.287.5459.1770

9. Gossner MM, Lewinsohn TM, Kahl T, Grassein F, Boch S, Prati D, et al. Land-use intensification causes multitrophic homogenization of grassland communities. Nature. Nature Research; 2016;540: 266–269. doi:10.1038/nature20575

10. Koerner SE, Smith MD, Burkepile DE, Hanan NP, Avolio ML, Collins SL, et al. Change in dominance determines herbivore effects on plant biodiversity. Nature Ecology and Evolution. 2018;2: 1925–1932. doi:10.1038/s41559-018-0696-y

11. Kohler AF, Gillet F, Gobat J, Buttler A. Seasonal vegetation changes in mountain pastures due to simulated effects of cattle grazing. Journal of Vegetation Science. 2004;15: 143–150.

12. Whitmore A. Impact of Livestock on Soil. In: Hartung J, Wathes CM, editors. Livestock Farming and the Environment. Hannover; 2001. pp. 39–41.

13. Olff H, Ritchie ME. Effects of herbivores on grassland plant diversity. Trends in Ecology and Evolution. 1998;13: 261–265. doi:10.1016/S0169-5347(98)01364-0

14. Marion B, Bonis A, Bouzillé J-B. How much does grazing-induced heterogeneity impact plant diversity in wet grasslands? Écoscience. 2010;17: 229–239. doi:10.2980/17-3-3315

15. Veldhuis MP, Howison RA, Fokkema RW, Tielens E, Olff H. A novel mechanism for grazing lawn formation: large herbivore-induced modification of the plant-soil water balance. Journal of Ecology. 2014;102: 1506–1517. doi:10.1111/1365-2745.12322

16. Hobbs RJ. Synergisms among habitat fragmentation, livestock grazing, and biotic invasion in Southwestern Australia. Conservation Biology. 2001;15: 1522–1528.

17. Bardgett RD, Wardle DA. Herbivore-mediated linkages between aboveground and belowground communities. Ecology. 2003;84: 2258–2268. doi:10.1890/02-0274

18. Breland TA, Hansen S. Nitrogen mineralization and microbial biomass as affected by soil compaction. Soil Biology and Biochemistry. 1996;28: 655–663.

19. Eisenhauer N, Milcu A, Nitschke N, Sabais ACW, Scherber C, Scheu S. Earthworm and belowground competition effects on plant productivity in a plant diversity gradient. Oecologia. 2009;161: 291–301. doi:10.1007/s00442-009-1374-1

20. Adler PB, Lauenroth WK. Livestock exclusion increases the spatial heterogeneity of vegetation in Colorado shortgrass steppe. Applied Vegetation Science. 2000;3: 213–222. doi:10.2307/1479000

21. Milchunas DG, Lauenroth WK, Burke IC. Livestock Grazing: Animal and Plant Biodiversity of Shortgrass Steppe and the Relationship to Ecosystem Function. Oikos. 1998;83: 65–74.

22. Ludwing J, Wilcox BP, Breshears DD, Tongway DJ, Imeson AC. Vegetation patches and runoff-erosion as interacting ecohydrological processes in semiaris landscapes. Ecology. 2005;86: 288–297. doi:10.1890/03-0569

23. Augustine DJ, Booth DT, Cox SE, Derner JD. Grazing intensity and spatial heterogeneity in bare soil in a grazing-resistant grassland. Rangeland Ecology and Management. 2012;65: 39–46. doi:10.2111/REM-D-11-00005.1

24. Kanivets S. The Factors and Conditions of Soil Formation: A Critical Analysis of Equivalence. Soil Science Working for a Living. 2017. pp. 3–8.

25. Laliberte E, Zemunik G, Turner BL. Environmental filtering explains variation in plant diversity along resource gradients. Science. 2014;345: 1602–1605. doi:10.1126/Science.1256330

26. Mráz P, Ronikier M. Biogeography of the Carpathians: evolutionary and spatial facets of biodiversity. Biological Journal of the Linnean Society. 2016;119: 528–559. doi:10.1111/bij.12918

27. Wypych A, Ustrnul Z, Schmatz DR. Long-term variability of air temperature and precipitation conditions in the Polish Carpathians. Journal of Mountain Science. 2018;15: 237–253. doi:10.1007/s11629-017-4374-3

28. Bakker ES, Ritchie ME, Olff H, Milchunas DG, Knops JMH. Herbivore impact on grassland plant diversity depends on habitat productivity and herbivore size. Ecology Letters. 2006;9: 780–788. doi:10.1111/j.1461-0248.2006.00925.x

29. Grace JB, Jutila H. The Relationship between Species Density and Community Biomass in Grazed and Ungrazed Coastal Meadows. Oikos. 1999;85: 398–408.

30. Borer ET, Seabloom EW, Gruner DS, Harpole WS, Hillebrand H, Lind EM, et al. Herbivores and nutrients control grassland plant diversity via light limitation. Nature. 2014;508: 517–520. doi:10.1038/nature13144

31. van Klink R, Schrama M, Nolte S, Bakker JP, WallisDeVries MF, Berg MP. Defoliation and Soil Compaction Jointly Drive Large-Herbivore Grazing Effects on Plants and Soil Arthropods on Clay Soil. Ecosystems. 2015;18: 671–685. doi:10.1007/s10021-015-9855-z

32. Grace JB, Anderson TM, Seabloom EW, Borer ET, Adler PB, Stanley Harpole W, et al. Integrative modelling reveals mechanisms linking productivity and plant species richness. Nature. 2016;529: 390–393. doi:10.1038/nature16524

33. Rudenko L, editor. National Atlas of Ukraine [Internet]. Kiev, Ukraine: Institute of Geography of National Academy of Sciences, Intelligence Systems Geo Ltd., and Ukrainian branch of World data center in Kiev Polytechnic Institute; 2007. Available: http://wdc.org.ua/atlas/en/

34. Havilah EJ. Forages and pastures: annual forage and pasture crops - species and varieties. Encyclopedia of Dairy Sciences. Second Edi. 2011. pp. 552–562. doi:10.1016/B978-0-12-374407-4.00193-X

35. Braun-Blanquet J. Plant sociology. The study of plant communities.[Translated, revised and edited by G.D. Fuller and H.S. Conrad]. First edit. New York and London: McGraw-Hill Book Co., Inc., New York and London; 1932.

36. R Core Team. R: A language and environment for statistical computing. R Foundation for Statistical Computing, Vienna, Austria. [Internet]. Vienna, Austria: R Foundation for Statistical Computing; 2017. p. https://www.r-project.org/. Available: https://www.r-project.org/

37. Rosseel Y. lavaan: An R Package for Structural Equation Modeling. Journal of Statistical Software. 2012;48: 1–36. doi:10.18637/jss.v048.i02

38. Grace JB. Structural equation modeling and natural systems. New York: Cambridge University Press; 2006.

39. Socher SA, Prati D, Boch S, Müller J, Klaus VH, Hölzel N, et al. Direct and productivity-mediated indirect effects of fertilization, mowing and grazing on grassland species richness. Journal of Ecology. 2012;100: 1391–1399. doi:10.1111/j.1365-2745.2012.02020.x

40. Hart RH, Samuel MJ, Test PS, Smith MA, Hart RH, Samuel MJ, et al. Cattle, Vegetation, and Economic Responses to Grazing Systems and Grazing Pressure. Journal of Range Management. 1988;41: 282–286.

41. Phelan P, Moloney AP, McGeough EJ, Humphreys J, Bertilsson J, O’Riordan EG, et al. Forage legumes for grazing and conserving in ruminant production systems. Critical Reviews in Plant Sciences. 2015;34: 281–326. doi:10.1080/07352689.2014.898455

42. Rutter SM. Diet preference for grass and legumes in free-ranging domestic sheep and cattle: Current theory and future application. Applied Animal Behaviour Science. 2006;97: 17–35. doi:10.1016/j.applanim.2005.11.016

43. Menneer JC, Ledgard S, Mclay C, Silvester W. The effect of a single application of cow urine on annual N2 fixation under varying simulated grazing intensity, as measured by four N15 isotope techniques. Plant and Soil. 2003;254: 469–470.

44. Grime JP. Competitive exclusion in herbaceous vegetation. Nature. 1973;242: 344–347. doi:10.1038/242344a0

45. Tardella FM, Catorci A. Context-dependent effects of abandonment vs. grazing on functional composition and diversity of sub-Mediterranean grasslands. Community Ecology. 2015;16: 254–266. doi:10.1556/168.2015.16.2.13

46. Symstad AJ. A Test of the Effects of Functional Group Richness and Composition on Grassland Invasibility. Ecology. 2000;81: 99–109.

47. Landsberg J, James CD, Maconochie J, Nicholls AO, Stol J, Tynan R. Scale-related effects of grazing on native plant communities in an arid rangeland region of South Australia. Journal of Applied Ecology. 2002;39: 427–444. doi:10.1046/j.1365-2664.2002.00719.x

48. Tzialla CE, Veresoglou DS, Papakosta D, Mamolos AP. Changes in soil characteristics and plant species composition along a moisture gradient in a Mediterranean pasture. Journal of environmental management. 2006;80: 90–98.

49. Grace JB, Pugesek BH. A Structural Equation Model of Plant Species Richness and Its Application to a Coastal Wetland. The American Naturalist. 1997;149: 436–460.

50. Carmel Y, Kadmon R. Effects of grazing and topography on long-term vegetation Mediterranean ecosystem in Israel. Plant Ecology. 1999;145: 243–254. doi:10.1023/A:1009872306093

51. Kholiavchuk D, Cebulska M. The highest monthly precipitation in the area of the Ukrainian and the Polish Carpathian Mountains in the period from 1984 to 2013. Theoretical and Applied Climatology. Theoretical and Applied Climatology; 2019;138: 1615–1628. doi:10.1007/s00704-019-02910-z

52. Buzhdygan OY, Meyer ST, Weisser WW, Eisenhauer N, Ebeling A, Borrett SR, et al. Biodiversity increases multitrophic energy use efficiency, flow and storage in grasslands. Nature Ecology & Evolution. 2020;4: 393–405. doi:10.1038/s41559-020-1123-8

53. Scherber C, Eisenhauer N, Weisser WW, Schmid B, Voigt W, Fischer M, et al. Bottom-up effects of plant diversity on multitrophic interactions in a biodiversity experiment. Nature. Nature Publishing Group, a division of Macmillan Publishers Limited. All Rights Reserved.; 2010;468: 553–556. doi:10.1038/nature09492

54. Ebeling A, Rzanny M, Lange M, Eisenhauer N, Hertzog LR, Meyer ST, et al. Plant diversity induces shifts in the functional structure and diversity across trophic levels. Oikos. 2018;127: 208–219. doi:10.1111/oik.04210

55. Ebeling A, Meyer ST, Abbas M, Eisenhauer N, Hillebrand H, Lange M, et al. Plant diversity impacts decomposition and herbivory via changes in aboveground arthropods. PloS one. Public Library of Science; 2014;9: 1–8. doi:10.1371/journal.pone.0106529

56. Hertzog LR, Ebeling A, Weisser WW, Meyer ST. Plant diversity increases predation by ground-dwelling invertebrate predators. Ecosphere. 2017;8: 1–14. doi:e01990. 10.1002/ecs2.1990

57. Greenwood KL, McKenzie BM. Grazing effects on soil physical properties and the consequences for pastures: a review. Australian Journal of Experimental Agriculture. 2001;41: 1231–1250.

58. Baur B, Cremene C, Groza G, Rakosy L, Schileyko AA, Baur A, et al. Effects of abandonment of subalpine hay meadows on plant and invertebrate diversity in Transylvania, Romania. Biological Conservation. 2006;132: 261–273. doi:10.1016/j.biocon.2006.04.018

59. Courtesy of the Integration and Application Network, University of Maryland Center for Environmental Science [Internet]. Available: ian.umces.edu/symbols

